# Early Tracheal and Salivary miRNAs in Extremely Preterm Infants Predict BPD-related Pulmonary Hypertension

**DOI:** 10.64898/2026.06.17.732493

**Authors:** Tang Li, Shaoyi Zhang, Vincent Aluquin, Ann Donnelly, Heather Stephens, Shakshi Sharma, Steven D Hicks, Dajiang Liu, Eric Austin, Roopa Siddaiah

## Abstract

Pulmonary hypertension (BPD-PH) associated with bronchopulmonary dysplasia (BPD) in preterm infants associates with high morbidity and mortality within the first two years of life. In a previous unbiased study, we identified a panel miRNAs in tracheal aspirates (TA) that were differentially expressed in extremely low gestational age newborns (ELGANs) with BPD-PH compared to those with BPD but no PH. To explore the predictive potential of these miRNAs, we studied TA exosomes from 7 days old ELGANs and analysed a curated panel of 16 miRNAs through logistic regression and calculated the predictive AUROC to diagnose BPD-PH at 36 weeks PMA. AUROC of TA miRNAs was 0.76 with sensitivity and specificity of 53% and 93%, respectively. Adding sex and gestational age to the variables improved the AUROC to 0.78 with sensitivity and specificity of 61 and 87% respectively. Due to challenges of obtaining TA in non-invasively ventilated infants, we collected saliva samples from ELGANs at 7 days of age and compared the log expression of these 16 miRNAs in both biofluids and found significant correlation in their expression (pearson r=0.92, p<0.001). We calculated the predictive AUROC of the same miRNAs to diagnose BPD-PH at 36 weeks PMA. AUROC of these miRNAs in saliva was = 0.85 with sensitivity and specificity of 82% and 72%, respectively; addition of biological sex and gestational age improved AUROC to 0.86 with sensitivity and specificity of 79% and 76% respectively. Leave-one-sample-out sensitivity analysis demonstrated stable training performance with reduced performance in testing samples, supporting the need for validation in larger independent cohorts. In conclusion, early salivary miRNAs have great potential for risk stratification of ELGANs to develop BPD-PH, while also providing the opportunity to identify target molecules and mechanisms that modulate molecular function.

## Introduction

Pulmonary hypertension (BPD-PH) associated with bronchopulmonary dysplasia (BPD) in preterm infants is associated with high morbidity and mortality within the first two years of life(1). A culmination of antenatal, perinatal, and postnatal injuries during the late canalicular stage disrupts developmental alveologenesis and angiogenesis, resulting in dysmorphic and reduced density of pulmonary vasculature(2). While impaired signaling in vascular growth factors plays a crucial role, a knowledge gap exists on the intricate relationship of genetics, epigenetics, inflammation, and transcription factors. Identification of the key drivers of BPD-PH early on will provide an opportunity to further study the underlying target pathways potential target molecule for preventative and therapeutic interventions.

In a previous study, to endotype molecular thumbprint of BPD-PH, we conducted an unbiased multiomics approach and identified a panel of miRNAs, mRNAs, and proteins in tracheal aspirates (TA) of preterm infants (3). We were particularly interested in the microRNAs which are small non-coding RNAs that regulate gene expression through intercellular signaling. While they play a crucial role inside the cell, a big proportion migrates outside (circulating miRNAs) bound to protein and through extracellular vesicles and be seen in bodily fluids such as blood, urine, saliva, breast milk, seminal fluid and tracheal aspirate (4, 5). Use of multimarker approach with multiple miRNA panels are gaining traction as a non-invasive method for diagnosis and prediction of disease progressions such as renal fibrosis in systemic lupus erythematosis and breast cancer(6, 7).

Because miRNAs are key regulators of lung organogenesis across both early and late developmental stages, our analysis focused on a panel of a panel of candidate miRNAs identified from prior studies and literature. We hypothesized that these miRNAs are expressed early in life in TAs and can predict development of BPD-PH at 36 weeks PMA and hence could help risk-stratify extremely low gestational age newborns (<28 weeks GA) (ELGANs) to optimize clinical management. Based on our previous study and literature review, we curated a panel of 16 miRNAs that are biological relevant to pulmonary vascular remodeling in developing lung namely hsa-let-7i-3p; hsa-let-7i-5p; hsa-miR-101-3p; hsa-miR-101-5p; hsa-miR-1255b-5p; hsa-miR-128-3p; hsa-miR-183-5p; hsa-miR-205-5p; hsa-miR-24-3p; hsa-miR-29a-3p; hsa-miR-3128; hsa-miR-3131; hsa-miR-501-3p; hsa-miR-542-3p; hsa-miR-624-5p; and hsa-miR-628-3p (3, 8–10).

Due to the inability to obtain TAs in ELGANs that are not intubated and on mechanical ventilation support, we also postulated that these miRNAs could be identified in saliva, which is a more readily accessible noninvasive biofluid. In a separate pilot study, we compared TA and saliva from ELGANs for overlapping miRNA profiles(11) and found a significant correlation between the salivary miRNAs and TA exosomal miRNAs. Given this result, we further hypothesized that early salivary expression of the panel of miRNAs can predict development of BPD-PH at 36 weeks PMA. We then used an in-silico model to explore the target gene pathways to understand the biological plausibility of the candidate miRNAs.

## Methods

We identified ELGANs that were born at <29 weeks of gestation admitted to the Penn State Health neonatal ICU. We excluded those with cyanotic cardiac defects or chromosomal abnormalities. Institutional informed consent was obtained from parents or guardians (IRB# STUDY00000482, Pennsylvania State University). TAs were collected during routine suctioning from ELGANs who were on mechanical ventilation support, within 7 days of age and stored at -80°C. We also collected saliva samples from all ELGANs irrespective of which respiratory support they were on. DNA genotek saliva collection kit (ORA-100) was used to collect saliva from ELGANs within first 7-days of age by placing the swab in the mouth for 60 seconds. The samples were stored at -80°C until further analysis. ELGANS were observed propectively. At 36 weeks PMA, routine echocardiograms was performed to screen for BPD-PH using parameters previously described(3). Echocardiogram features to suggest elevated right sided pressures such as tricuspid regurgitation jetvelocity, interventricular septal flattening, annular plane systolic excursion and pulmonary artery acceleration time were used to define BPD-PH.

### TA processing: Exosomal RNA extraction

Frozen tracheal aspirate samples were thawed on ice, and mixed by pipetting. For each sample, 300 uL of the aspirate was transferred to a 1.5 ml tube and 75 ul of ExoQuick Exosome Precipitation Solution (System biosciences, cat # EXOQ5A-1) was added. Sample and solution were mixed well by inverting, incubated upright and undisturbed at 4 C overnight. Sample mixture was then centrifuged at 1550 x g for 30 min at 4 C. After centrifugation, supernatant was discarded without disturbing the pellet. Pellet was then resuspended in 100 uL 1x PBS. All 100 uL of the sample was used for RNA extraction. RNA was extracted using Zymo Research Quick-RNA Micro Prep Kit following manufacturer’s protocol. The typical size distribution for the small RNA ranged between 13.14 and 283.50 ng.

### Salivary samples processing

Frozen saliva samples were thawed on ice. RNA was extracted using Quick-RNA Microprep purification kits (Zymo Research). RNA Pico BioAnalyzer (Agilent technologies) was used to determine the RNA quality and quantity. The typical size distribution for the small RNA and quantities for saliva ranged between 33.90 and 922.50 ng

### RNA sequencing for miRNA

Small RNA libraries were prepared from 5–250 ng total RNA using the QIAseq miRNA Library Kit (QIAGEN) as per the manufacturer’s instructions. The built-in Unique Molecular Identifier (UMI) application used to eliminate possible PCR duplicates in sequencing datasets to facilitate unbiased gene expression profiling. The unique barcode sequences were then incorporated in the adaptors for multiplexed high-throughput sequencing. The final product was assessed for its size distribution and concentration using a BioAnalyzer High Sensitivity DNA Kit (Agilent Technologies). Libraries were prepared in the Penn State College of Medicine Genome Sciences core (RRID:SCR_021123).

The libraries were pooled and diluted to 3 nM using 10 mM Tris-HCl, pH 8.5 and denatured. The denatured libraries were then loaded onto an SP flow cell on an Illumina^®^ NovaSeq 6000 instrument and run for 68 cycles according to the manufacturer’s instructions. De-multiplexed sequencing reads were generated using Illumina bcl2 fastq (released version 2.20.0.422), allowing no mismatches in the index read. Fitlered reads were aligned to the human reference genome (GRCh38 using HISAT2 (version 2.1.0)(12) and the resulting ded-duplicated reads were summarized to each gene using HTSeq(13). All RNA seq experiments were conducted at the Penn State College of Medicine Genome Sciences Core Facility. Data sets for both TA and saliva samples were uploaded to GEO https://www.ncbi.nlm.nih.gov/geo/query/acc.cgi?acc=GSE326620

### Data analysis

Statistical analysis was performed with R (version 4.3.1). Differential expression analysis was performed using edgeR to evaluate miRNAs associated with disease status in both saliva and tracheal aspirate samples. In multivariable logistic regression models, raw counts from the 16 candidate miRNAs were used as predictors. Models were fitted with and without adjustment for gestational age (in weeks) and biological sex. Finally a leave-one-sample-out-sensitivity analysis was conducted by removing one sample at a time, refitting the model on the remaining samples and recalculating model performance on the reduced dataset. The resulting sample-specific accuracies were plotted. To correlate miRNA expression across both biofluids, we compared the log expression levels of the candidate 16 miRNA in both TA and saliva.

### Pathway analysis

To identify the predicted target genes for the panel of miRNAs, we used Ingenuity Pathway Anlaysis (IPA software, Qiagen). The core function analysis was performed for to identify molecular functions based on prediction scores and direct and indirect relationships using Ingenuity knowledge base.

## Results

We consented and recruited 104 subjects and scheduled sample collections for days 3 and 7 after birth; however, samples were obtained for 58 subjects within the first week of life due to timing of the consent obtained. In total we were able to procure 29 TA samples and 54 saliva samples (Figure 1). Eight of these 58 subjects were on room air at 36 weeks gestation and were not included in the study. Of the remaining 50 subjects who obtained a screening echocardiogram at 36 weeks, 21 showed the above mentioned criteria for PH and were categorized into BPD-PH. There was no significant difference in demographics in the two groups in terms of their gestational age, birth weight, sex and other key perinatal factors (Table 1). Of the 21 subjects, 4 of them died before 36 weeks PMA, and because they had eachocardiographic findings of PH and CXR consistent with BPD changes, they were included in the BPD-PH cohort.. Of the 29 TA samples, 16 were from ELGANs with BPD and 13 from ELGANs with BPD-PH. Of the 54 saliva samples, 25 were from ELGANs with BPD and 29 from ELGANs with BPD-PH.

**Figure 1:**
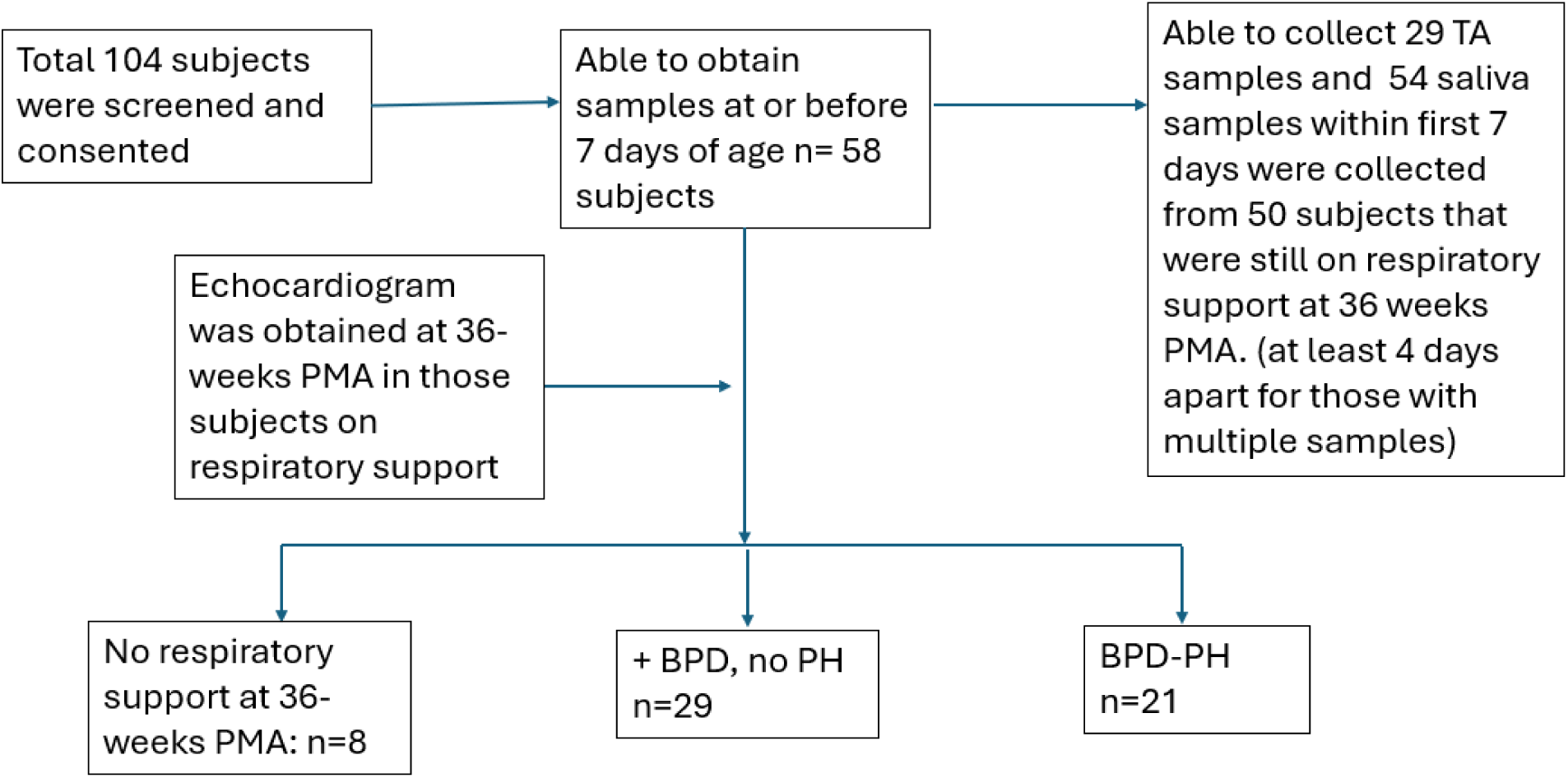
Flow diagram illustrating study design.

A multivariate logistic regression model was fitted for the 16 candidate miRNAs as predictors from TAs to predict BPD-PH (BPD=16, BPD-PH=13) at 36 weeks PMA. The unadjusted model showed an AUROC of = 0.75 with sensitivity and specificity of 54% and 93%, respectively (Figure 2A). After adjusting for gestaional age and the sex into the regression modeling, there was an improvement in the AUROC = 0.78 with sensitivity and specificity of 62% and 88% respectively (Figure 2B). To further evaluate model robustness, we conducted a leave-one-sample-out sensitivity analysis, which demonstrated stable training classification accuracy above 70% (Figure 2c). Testing accuracy results are provided in Supplementary-Figure 1A. Logistic regression summaries for both the null and adjusted models including sex and gestational age are provided in Supplementary-Table 1.

**Figure 2.**
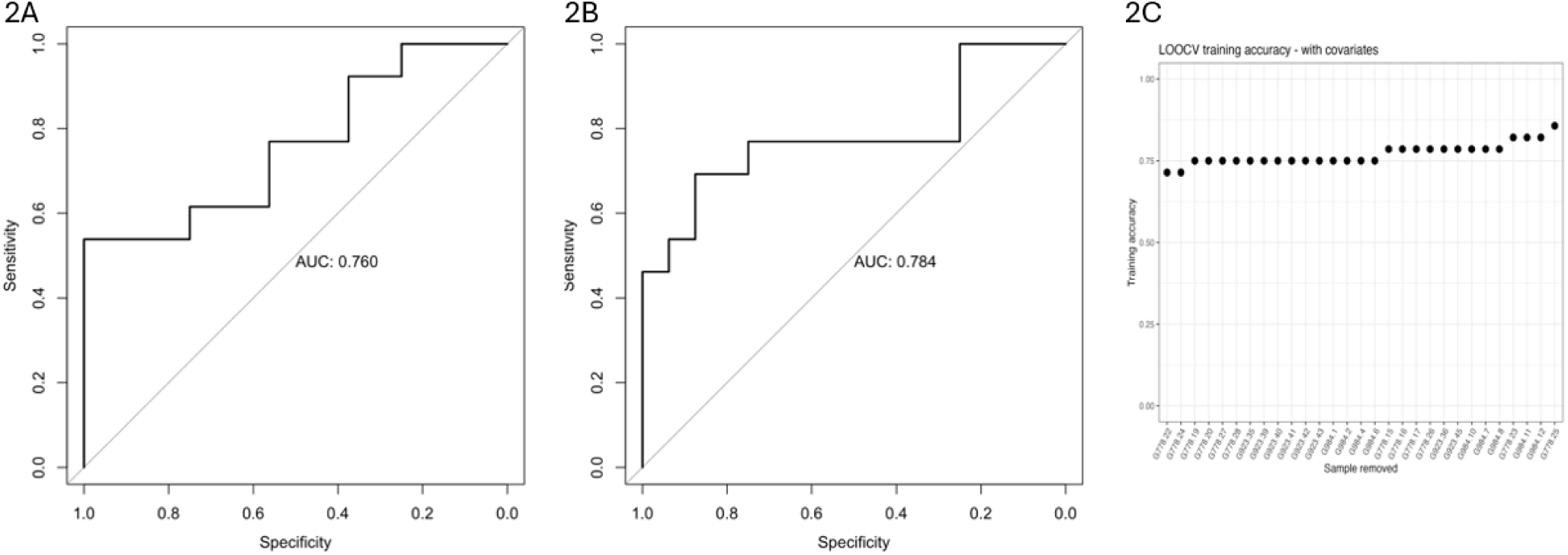
Figure 2A: Receiver Operating Curve (ROC) for predicting development of BPD-PH based on candidate miRNA panel in TA collected at 7 days of age. Figure 2B: Receiver Operating Curve (ROC) for predicting development of BPD-PH based on candidate miRNA panel in TA collected at 7 days of age on adjusting for gestational age and sex. Figure 2C: Leave-one-sample-out-sensitivity analysis showing >90% accuracy.

A correlation analysis of expression of 16 miRNAs in both saliva and TA revealed a significant positive correlation in log expression (Pearson r=0.92, p<0.001) (n=50) (Figure 3). A multivariate logistic regression model was fitted for 16 candidate miRNAs in salivary samples to predict BPD-PH (BPD= 28, BPD-PH=23) at 36 weeks PMA. The unadjusted model showed an AUROC of = 0.85 with sensitivity and specificity of 82% and 72% respectively (Figure 4A). On adding gestational and sex to the logistic regression model we again saw an improvement in the AUROC = 0.86 with sensitivity and specificity of 79% and 76% respectively (Figure 4B). Leave-one-sample-out sensitivity analysis demonstrated stable training classification accuracy above 75% (Figure 4c), while testing accuracy results are shown in Supplementary-Figure 1B. Logistic regression summaries for both the null and adjusted models including sex and gestational age are provided in Supplementary-Table 2.

**Figure 3:**
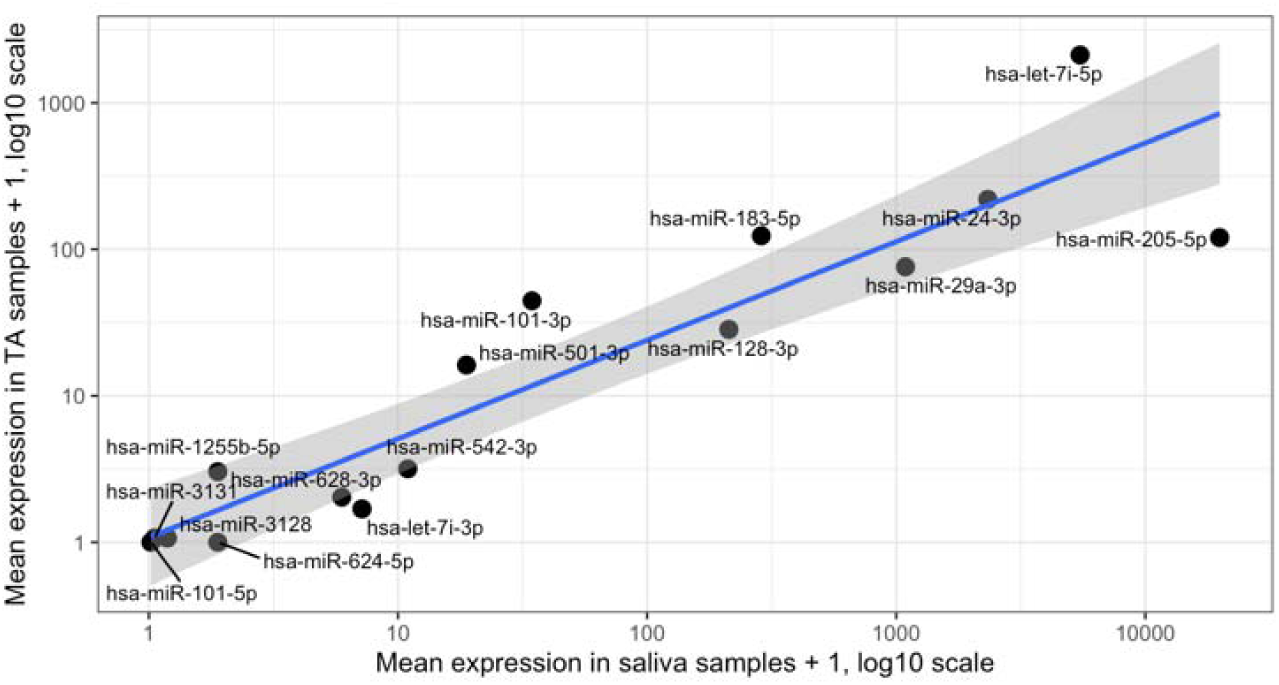
A correlation analysis of the log mean expression from 16 miRNAs in saliva and tracheal aspirate showed significant positive correlation r=0.92, p<0.001

**Figure 4.**
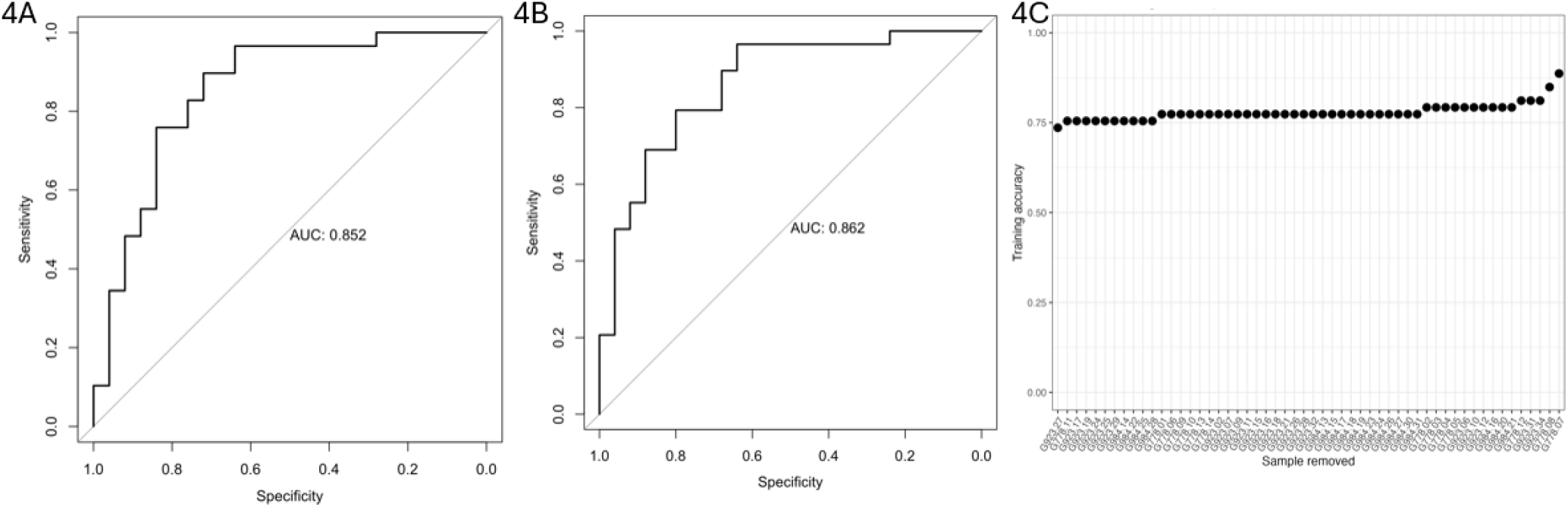
Figure 4A: Receiver Operating Curve (ROC) for predicting development of BPD-PH based on candidate miRNA panel in saliva collected at 7 days of age. Figure 4B: Receiver Operating Curve (ROC) for predicting development of BPD-PH based on candidate miRNA panel in salvia collected at 7 days of age on adjusting for gestational age and sex. Figure 4C: Leave-one-sample-out-ssensitivity analysis showing >80% accuracy.

Using an in-silico Inguinty Pathway Analysis, we identified key target genes that were involved in the pathways of these target miRNAs. Some of the key genes we identified were PDGFA, TNF, ANGTL4 and PTGS2 and the specific miRNA in the panel miR29a seemed to play and important role in the interaction of these genes (Figure 5). Of the identified panel of miRNAs, miR29a was of particular interest and most significantly overexpressed in infants with BPD-PH. This miRNA is shown to target TGF-β signaling during lung development and specifically affect differentiation of smooth muscle cells, disrupting development of distal lung vasculature(14, 15).

**Figure 5.**
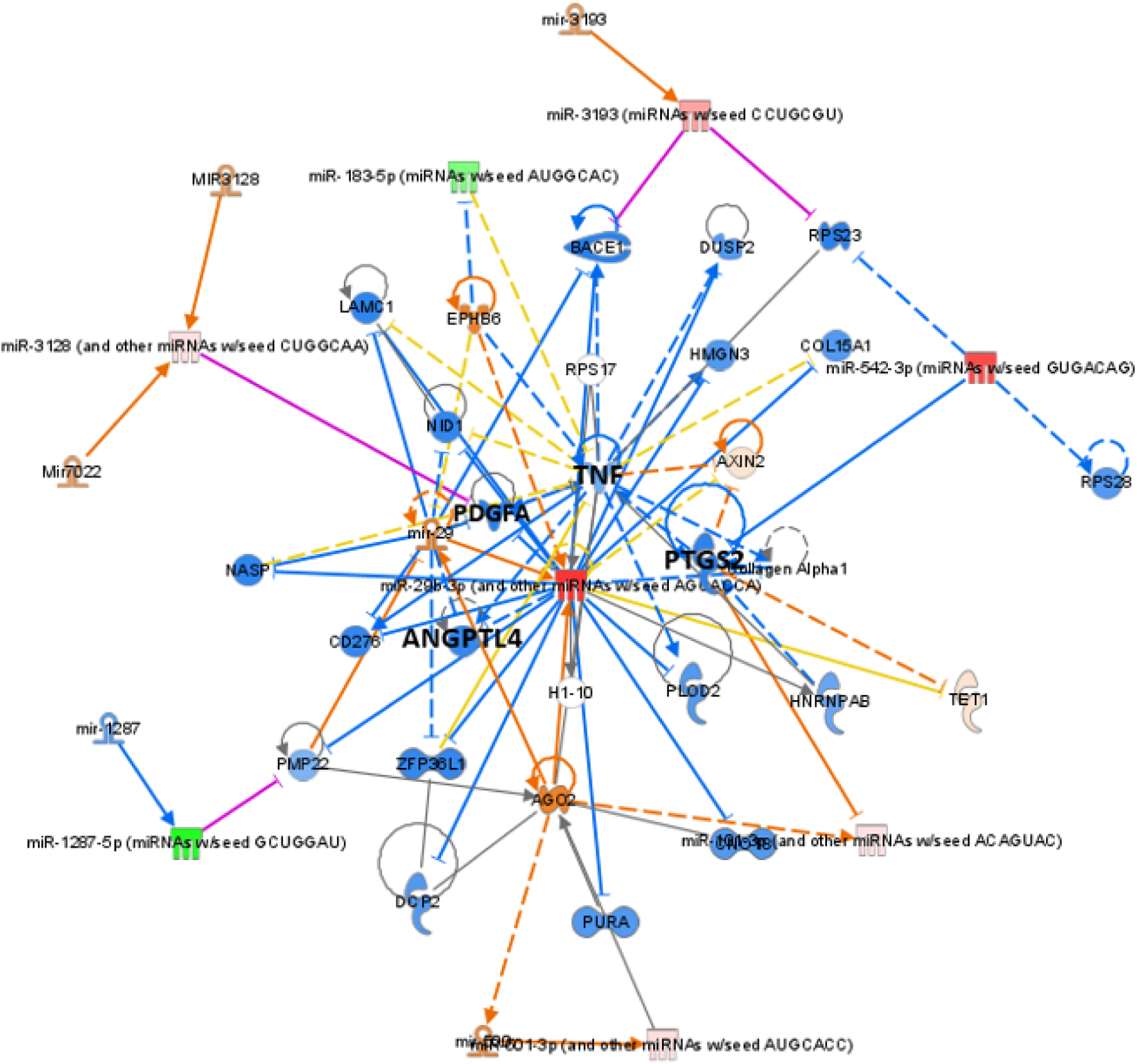
IPA pathways: Inguinity pathway analysis of the candidate miRNA panel.

## Discussion

Noninvasive biomarkers play an important role in studying pathology, especially in vulnerable populations such as ELGANS where noxious stimuli lead to poorer outcomes. In our pilot study, we identified a panel of miRNAs upregulated in BPD-PH; however, TA collection is limited to infants who are on mechanical ventilation, which skews our study population to sicker infants and is impractical for studying longitudinal changes. In this study, we not only identified early TA miRNAs showed strong classification performance within the study dataset, we also replicated this in early salivary samples in ELGANs. Thus, saliva may serve as an alternate and accessible biofluid in ELGANs to study miRNAs in lung development and maturation. It also allows for longitudinal monitoring of lung maturation both during initial NICU stay and after discharge to outpatient care.

Given the complex pathobiology of BPD and BPD-PH, we believe that a multimarker approach rather than a single biomarker is more appropriate for prediction and prognostic purposes. Especially in vulnerable population such as preterm infants who sustain physiological changes secondary to noxious stimuli such as blood draws, a non-invasive and painless approach is desired. Previous work highlighted that features of PH by echocardiogram at day 7 of life could predict BPD-PH(16). However, echocardiogram’s low negative predictive value may result in high false positives and hence increased frequency of echocardiogram than what is necessary for close monitoring of these infants. Combining echocardiographic imaging with molecular biomarkers could potentially improve early risk stratification of preterm infants. Incorporating gestational age and sex into the predictive model significantly enhanced its discriminatory performance as indicated by a higher AUROC. This finding aligns with established pathophysiology where lower gestational age is independently associated with wn increased incidence and severity of BPD and BPD-PH due to premature exposure of developing lung to mechanical and hyperoxia stress during critical phases of alveolarization and vascular growth.

Tracheal aspirates (TA) are enriched with extracellular vecsicles which play a key role in lung aveolar epithelial cells and endothelial cells as potentially crucial contributors to lung growth and development. In addition, TA may serve as a suitable liquid biopsy of the lungs (17). In our previous study, we found a significant overlap in the tracheal aspirate exosomal and salivary miRNA profile (11). Systemic release of intracellualr miRNAs facilitated by either vesicular transportation or protein binding, enables their detection in various biofluids. This mechanism likely contributes to the observed concordance in miRNA expression profiles between TA exosomes and saliva. Moreover, Identifying the target gene expression using in-silico modeling (Inguinity Pathway Analysis), we identified key markers such as PDGFA, ANGPTL4 and TNF which have been shown to play a key role in angiogenesis which is crucial for lung development(18–21).

Many miRNAs in our panel have been studied extensively to modulate genes responsible for lung organogenesis and growth and development. One of our highly differentially expressed miRNAs, miR29a has been shown to be over-expressed in hyperoxia-induced BPD in mouse models, and its inhibition has been shown to alleviate lung injury(22). Further, disruption of miR29a in-vivo leads to immature vascular smooth muscle cell differentiation and altered phenotype in distal lung vasculature(14). Given the key role of vascular smooth muscle differentiation, proliferation, and migration in pulmonary vascular development during lung organogenesis, it is crucial to delineate the key pathways. Because experimental PDGFA deletion induces lung development arrest(23), we postulate that miR29a plays a key role by stunting alveolo-vasculogenesis through decreased expression of PDGFA. Similarly, miR101-3p was found to suppress High-mobility Group Protein 3 (HMGB3) and TGF-β1/Smad3 signaling pathways which are involved in inflammation and fibrotic remodeling in BPD mouse model (24). Upregulation of miR-let-7 was associated with modulation of TGF--β1 signaling pathway mediated by Lipoxin A in hyperoxia murine BPD model(25).

Our study is strengthened by further interrogating a panel of miRNAs that were identified through an unbiased approach in our previous study for their predictive value. While our previous study demonstrated overlapping in overall miRNA profile in TAs and saliva, our current study findings show similar candidate miRNA expression in both biofluids. Notably, the predictive values of these miRNAs are highly similar across both sources. We were able to address some of the developmental stages related alterations in the miRNAs by adding gestational age and sex into our regression modeling which improved our predictive scores. We further explored the biological plausibility of the panel of miRNAs through an Ingenuity pathway analysis to identify key underlying pathways playing a role in pathobiology of BPD-PH.

Our study is limited as a single-center-study with relatively small number for predictive analysis of the miRNA and needs to be validated in a large multicenter study. We also screened for BPD-PH using echocardiogram since cardiac catheterization is highly invasive and less frequently used. However, catheterization is not generally employed as a screening tool for BPD-PH and to minimize inter-observer variability, all the echocardiograms were evaluated by a single cardiologist. With 29 TA and 54 saliva samples, modeling of 16 miRNAs introduces a potential risk of overfitting. While we utilized leave-one-sample out analysis to assess sensitivity of the data, external validation in an independent cohort is necessary to assess generalizability. Consistent with published finidings, Established literature identifies several clinical variables that pre-dispose preterm infants to BPD-PH, specifically: fetal growth restriction, maternal pre-eclampsia, chorioamnionitis and prolonged mechanical ventilation. However, given our sample size, these variables will be futher validated in a future larger cohort study.

In summary, our select panel of miRNAs in both TA and saliva from seven-day-old ELGANs predicted the development of BPD-PH. Salivary miRNAs are excellent noninvasive biomarkers for longitudinal follow-up, assamples can be conveniently collected at home or clinical setting after NICU. Risk stratification of preterm infants at risk to develop BPD-PH in the future could help guide clinical management with tighter monitoring of fluid balance, avoiding rapid weaning of respiratory support and addressing intracardiac shunting appropriately. In addition, further mechanistic studies of the miRNAs has the potential to identify target therapeutics for early intervention and preventative strategies.

## Acknowledgments

All protocols were performed in compliance with all relevant ethical regulations approved by Penn State Health IRB (STUDY 00000482); written informed consent was obtained from all patients who participated in this study. Diane Kitch for maintaining upto date regulatory documents of the study.

## Author Contribution

RS: Conceptulization, study design, data analysis, data interpretation, manuscript preparation

TL: Data analysis, editing manuscript

SZ: Data analysis, editing manuscript

VA: Study design, Data analysis, editing manuscript

AD: Screening, consenting, Sample collection, storage, editing manuscript

HS: Screening, consenting, Sample collection, editing manuscript

SS: Data management, editing manuscript

SH: Study design, data interpretation, editing manuscript

DL: Data analysis, editing manuscript

EA: Study design, data interpretation, editing manuscript

