## Supplementary Table 1 for "Early Tracheal and Salivary miRNAs in Extremely Preterm Infants Predict BPD-related Pulmonary Hypertension"

| model | term | estimate | std.error | statistic | p.value |
| --- | --- | --- | --- | --- | --- |
| full_GLM_without_covariates | (Intercept) | -0.3175366 | 0.54962243 | -0.5777359 | 0.56344244 |
|  | `hsa-let-7i-5p` | -0.0008205 | 0.00060141 | -1.3643217 | 0.17246632 |
|  | `hsa-miR-101-3p` | 0.00111361 | 0.00558382 | 0.19943478 | 0.84192265 |
|  | `hsa-miR-128-3p` | 0.018918 | 0.04007419 | 0.47207447 | 0.63687363 |
|  | `hsa-miR-183-5p` | 0.00289714 | 0.01358721 | 0.21322543 | 0.83115113 |
|  | `hsa-miR-205-5p` | 0.00118071 | 0.00456659 | 0.25855409 | 0.79597931 |
|  | `hsa-miR-24-3p` | -0.0019654 | 0.0027808 | -0.706773 | 0.47970754 |
|  | `hsa-miR-29a-3p` | 0.02167503 | 0.02052062 | 1.05625571 | 0.29085141 |
| full_GLM_with_covariates | (Intercept) | 6.27179371 | 9.12917603 | 0.68700545 | 0.49207929 |
|  | `hsa-let-7i-5p` | -0.000843 | 0.00064397 | -1.3090837 | 0.19050601 |
|  | `hsa-miR-101-3p` | 0.00194677 | 0.00655432 | 0.2970204 | 0.76645093 |
|  | `hsa-miR-128-3p` | 0.03393541 | 0.06024116 | 0.56332591 | 0.57321298 |
|  | `hsa-miR-183-5p` | 0.00023196 | 0.01896512 | 0.01223097 | 0.99024134 |
|  | `hsa-miR-205-5p` | 0.00191733 | 0.00554992 | 0.34546974 | 0.72974125 |
|  | `hsa-miR-24-3p` | -0.00193 | 0.00310221 | -0.6221511 | 0.53384248 |
|  | `hsa-miR-29a-3p` | 0.0222159 | 0.02819066 | 0.78805901 | 0.43066219 |
|  | GA_raw | -0.278439 | 0.35556803 | -0.7830822 | 0.43357884 |
|  | sex011 | 0.76828425 | 1.06016872 | 0.72468111 | 0.46864768 |

Supplementary Table 1: Logistic regression summary results for tracheal aspirate miRNA models without and with sex and gestational age covariates
