## Supplementary Table 2 for "Early Tracheal and Salivary miRNAs in Extremely Preterm Infants Predict BPD-related Pulmonary Hypertension"

| model | term | estimate | std.error | statistic | p.value |
| --- | --- | --- | --- | --- | --- |
| full_GLM_without_covariates | (Intercept) | 0.54184792 | 0.64441505 | 0.84083684 | 0.40043935 |
|  | `hsa-let-7i-3p` | -0.016738 | 0.07933822 | -0.21097 | 0.83291068 |
|  | `hsa-let-7i-5p` | 0.00011082 | 0.0001219 | 0.90906594 | 0.36331532 |
|  | `hsa-miR-101-3p` | 0.00334113 | 0.01061713 | 0.31469189 | 0.7529956 |
|  | `hsa-miR-128-3p` | -0.0055795 | 0.00469611 | -1.1881027 | 0.23479295 |
|  | `hsa-miR-183-5p` | -0.0022629 | 0.00257955 | -0.8772312 | 0.38036108 |
|  | `hsa-miR-205-5p` | -1.739E-05 | 5.5014E-05 | -0.3161726 | 0.75187153 |
|  | `hsa-miR-24-3p` | 0.00052516 | 0.00042517 | 1.23516252 | 0.21677002 |
|  | `hsa-miR-29a-3p` | -0.0009995 | 0.00093121 | -1.0732957 | 0.28313848 |
|  | `hsa-miR-501-3p` | 0.06074597 | 0.03089445 | 1.96624228 | 0.04927063 |
|  | `hsa-miR-542-3p` | 0.06903284 | 0.06037782 | 1.14334771 | 0.25289425 |
|  | `hsa-miR-628-3p` | -0.1586369 | 0.09830903 | -1.6136555 | 0.10660218 |
| full_GLM_with_covariates | (Intercept) | -7.2062686 | 7.70417946 | -0.9353713 | 0.34959697 |
|  | `hsa-let-7i-3p` | -0.0215918 | 0.0794575 | -0.27174 | 0.78582191 |
|  | `hsa-let-7i-5p` | 0.00010351 | 0.00012136 | 0.85292412 | 0.39370138 |
|  | `hsa-miR-101-3p` | 0.00065502 | 0.01071127 | 0.06115243 | 0.95123781 |
|  | `hsa-miR-128-3p` | -0.0056524 | 0.00481131 | -1.1748207 | 0.24006644 |
|  | `hsa-miR-183-5p` | -0.0015028 | 0.0023729 | -0.6333268 | 0.52652023 |
|  | `hsa-miR-205-5p` | -3.742E-05 | 5.7911E-05 | -0.6462108 | 0.51814282 |
|  | `hsa-miR-24-3p` | 0.00065731 | 0.0004402 | 1.49320029 | 0.13538477 |
|  | `hsa-miR-29a-3p` | -0.0011672 | 0.00095497 | -1.2222695 | 0.22160574 |
|  | `hsa-miR-501-3p` | 0.0676132 | 0.0324941 | 2.08078418 | 0.03745366 |
|  | `hsa-miR-542-3p` | 0.0891251 | 0.06343081 | 1.40507582 | 0.15999873 |
|  | `hsa-miR-628-3p` | -0.1631007 | 0.10199419 | -1.5991179 | 0.10979441 |
|  | GA_raw | 0.29321969 | 0.2968797 | 0.98767174 | 0.32331344 |
|  | sex01 | 0.36517573 | 0.71816095 | 0.50848731 | 0.61111164 |

Supplementary Table 2: Logistic regression summary results for salivary miRNA models without and with sex and gestational age covariates
